# Analysis of *E. coli* growth dynamics during Lambda(vir) phage infection reveals phage decay

**DOI:** 10.1101/2023.11.28.569039

**Authors:** Benjamin M. Gitai, Joseph P. Sheehan

## Abstract

Biological interactions like those between a pathogen and a host are intrinsically dynamic. The Luria-Delbruck experiment is a classic quantitative analysis demonstrating that the mutation rates of bacterial hosts are independent of the presence of their bacteriophage pathogens. However, this experiment was performed as an endpoint analysis. To determine if new insights could be gained from measuring the dynamics of bacterial-host interactions, we used a high-throughput spectrophotometer to repeat this classic experiment. We examined interactions between *Escherichia coli* bacteria and the lytic lambda(vir) phage to confirm the fundamental finding of the Luria-Delbruck experiment. Specifically, we demonstrated that the variance in the timing required for phage-resistant mutants to emerge increased as the size of the starting bacterial population decreased. In addition to confirming the Luria-Delbruck results, our analysis of the dynamics of bacterial growth in the presence of phage revealed unexpected findings, including the significant decay in lambda(vir) potency and the substantial impact of dead bacterial debris on culture density measurements. Phage are typically thought to be extremely stable, such that our discovery that their potency rapidly decays in experimental settings may have implications for real-world applications where phage potency is important, such as phage therapy pharmacology. Our results thus establish a more rapid method of analyzing bacterial resistant mutant fluctuations and illustrate how dynamical measurements can provide new insights into even extremely well-characterized experimental systems.

## Introduction

Nearly all biological systems involve dynamic interactions that change over time. As a result, studies that only measure endpoints and do not consider the dynamics across the experiment are susceptible to missing important features of the interactions. Just as a movie cannot be described simply by the first and last frames of the film, analyzing an experiment solely by endpoints misses out on dynamic information. In sports, the winning team is just the team with the higher score at the end of the game, but they may not be the better team, which is why advanced in-game metrics that track dynamics over the course of the game outperform final score as predicters [1]. Thus, measuring and analyzing information about interaction dynamics during an event or experiment are necessary to fully understand a system.

The Luria-Delbruck experiment is one of the most famous experiments in evolutionary biology, representing a classic demonstration that mutation is random and not induced by the environment [2]. Luria and Delbruck studied the interactions between bacteria and bacteriophage (phage), which are viruses that infect and kill bacteria. The bacteria can mutate to become resistant to the phage. To determine if the rate at which the bacteria mutated to become phage-resistant was constant or induced by the presence of the phage, Luria and Delbruck took advantage of a fundamental feature of statistical noise that the larger a sample size becomes, the less fluctuation there is in the mean of the sample. For example, if one were to flip a coin 1000 times, the outcome would tend to be closer to 50% heads than if one were to only flip a coin 10 times [3]. To implement this idea experimentally, Luria and Delbruck grew bacterial cultures of varying population sizes with phage to see if the percentage of bacteria that mutated and became resistant to the phage changed based on the size of the bacterial population. They found that the variance in the percentage of the phage-resistant bacteria decreased as the population size increased [2]. This trend is the same as for random events like coin flips, implying that mutation is random rather than induced.

One caveat to the Luria-Delbruck experiment was that they used an endpoint analysis [2], such that there might be aspects of the phage-bacteria interaction that were missed. When the Luria-Delbruck experiment was performed in 1943, high-throughput, multi-time point systems for measuring the dynamics of bacterial growth did not exist. Now that tools like automated 96-well plate spectrophotometers exist, one can measure bacterial growth dynamics throughout an entire experiment. These new instruments allow us to not only measure individual bacterial growth curves, but also measure variance, providing us with new insights into the dynamics of the Luria-Delbruck experiment. Here we demonstrate that by analyzing the dynamics, rather than just the endpoints, of phage-bacterial interactions, it is possible to quickly reproduce the conclusions of Luria-Delbruck. Additionally, we discovered new phenomena, including the unexpected degradation of phage potency during the course of the experiment.

## Results

### Growth dynamics reveal that the variance in lambda(vir) phage resistance depends on *E. coli* population size

In the classic experiment by Luria and Delbruck[2], the fluctuation in the number of *E. coli* resistant to a phage infection was shown to be much greater than the mean, demonstrating that the resistance mutations were random and pre-existed in the population, rather than being induced by the phage. We hypothesized that similar trends would occur in growth dynamics, as populations with more resistant mutants would result in earlier population-level growth in the presence of the phage. To test this hypothesis, we measured the dynamics of *E. coli* MG1655 growth in the presence of lambda(vir) phage. We chose to focus on lambda(vir) because it is exclusively lytic and thus reduces complications associated with the possibility of lysogeny [4].

In a typical *E. coli* lambda(vir) infection growth curve, one can observe several characteristic features (Figure 1A). We first observed an initial increase in optical density (OD). This likely reflects an initial increase in bacterial growth, because our experiments started with fewer phage than bacteria. We then saw a rapid decrease in OD, reflecting bacterial lysis by the phage once the number of phage exceeded that of the bacteria. Following this decrease in OD, the OD leveled out. The flattening of the growth curve is likely due to the fact that the number of surviving bacteria is below the limit of detection of the plate reader spectrophotometer. Later, we observed a second stage of growth in OD, likely caused by the surviving, phage-resistant, bacteria growing in population size. The time required for this second phase of growth to begin should be inversely proportional to the number of phage-resistant bacteria in a given culture because the more resistant bacteria present, the earlier this second growth phase should arise (Figure 1A).

**Figure 1.**
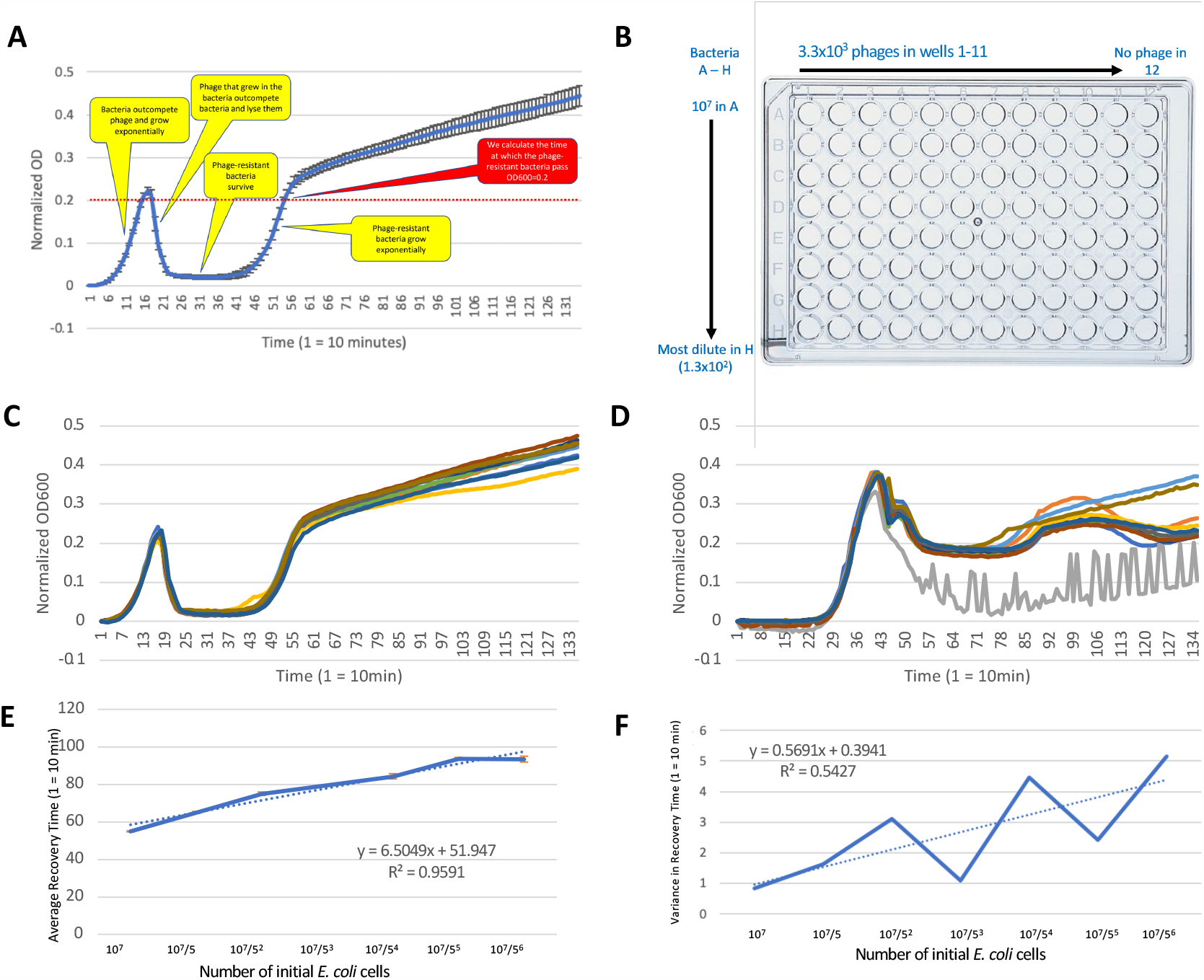
(A) Illustration of a typical growth curve from the experiment set up in (B), highlighting in yellow the different features of the growth curve: an initial phase where bacteria outcompete phages resulting in increased OD600, collapse when phage numbers overtake bacterial numbers, and recovery of the culture when phage-resistant bacteria subsequently grow. We also highlight in red the point we calculate as the recovery phase (the second time the culture passes OD = 0.2). (B) Schematic of the experimental setup for replicating the Luria-Delbruck experiment in a 96-well plate. For each row, columns 1-11 were replicates of the same experiment in which 3.3x10^3^ phages were mixed with the starting number of bacteria noted. No phages were included in column 12 as a control for bacterial growth. Each row represented a 5-fold dilution of bacteria, with 10^7^ bacteria in row A and 1.3x10^2^ bacteria in row H. (C) The growth curves for row A (starting with 10^7^ bacteria per well). (D) The growth curves for row E (starting with 10^7^/5^4^ bacteria per well). For C-D the colors represent arbitrary replicates. (E) The average time required for each dilution (experimental row) to recover past OD = 0.2. (F) The variance in the time required for each dilution to recover, with dotted line showing the best fit linear regression. We note that for E-F, we do not include row H because most of the wells did not recover.

Luria-Delbruck predicts that the variance of resistance frequency should be inversely proportional to the initial population size of *E. coli*. As a result, if growth dynamics can indeed be used to analyze population resistance frequencies, we would expect that reducing the initial numbers of *E. coli* should increase the variance in the time required for observing the second stage of bacterial growth. We thus performed the following experiment in a 96-well plate (Figure 1B): within each row we had positive and negative growth control wells in addition to 10 independent wells with the same *E. coli* starting population size and a fixed number of phage particles. Between each subsequent row, we diluted the number of starting bacteria 5-fold while keeping the number of phage constant. For example, each well in Row A had 1x10^7^ *E. coli*, while each well in Row H had 1.28 × 10^2^ *E. coli* (Figure 1B). All wells received the same number of lambda(vir) phage (3.3x10^3^ plaque forming units).

To quantify the time to the second growth phase, we measured the time point at which the OD600 of the culture exceeded 0.2 for the second time (dotted red line in Figure 1A). We found that as the population size decreased, the average time to exceed 0.2 increased. Representative growth curves from Row A and E are shown in Figure 1C and Figure 1D, respectively, and the averages across each row are shown in Figure 1E. This was expected, because starting at a lower population would mean that it takes more time to reach a given OD. We also found that the variance in time required to exceed an OD600 of 0.2 increased as population size decreased (Figure 1F). This trend is in agreement with the results of the Luria-Delbruck experiment, thereby validating that the use of growth dynamics is an effective way of measuring resistance rate fluctuations.

#### Lambda(vir) potency decays over time

When examining the growth curves measured in the initial experiment, we observed that as the initial population size decreased, the maximum OD reached by the population increased (Figure 2A). This was unexpected because the maximum OD should depend on when the number of phage in the population overtake the number of bacteria, and starting with fewer bacteria should change the timing of that process but not the relative numbers of bacteria and phages. We thus sought to understand the cause of the unexpected increase in maximum OD.

**Figure 2.**
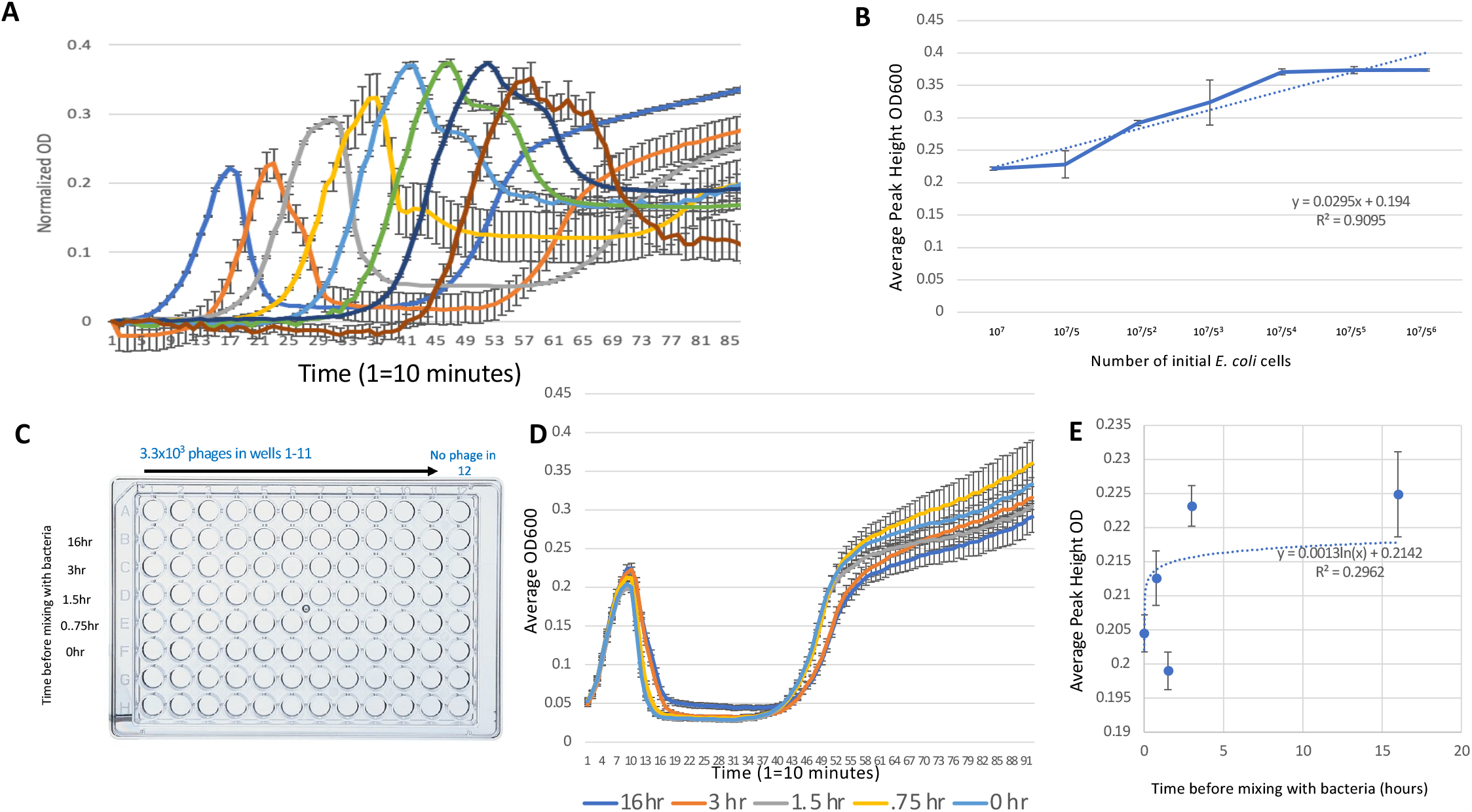
(A) Averages of the replicate wells from each row from the experiment in Figure 1 to focus on the heights of the initial OD peaks. Colors represent the rows illustrated in Figure 1D in descending order from most starting cells to fewest of: dark blue, orange, gray, yellow, light blue, green, navy, brown. (B) Average of the initial OD peak for each of the rows from Figure 1. (C) Schematic of the experimental setup to determine the effect of incubation with shaking at 37°C for different lengths of time on phage potency. For each row, columns 1-11 were replicates of the same experiment in which 3.3x10^3^ phages were mixed with the 10^7^ bacteria after the length of time indicated on the right for each row. (D) The average of the OD600 curves for each of the rows shown in C. Colors for each delay indicated below chart. (E) Average of the peak height OD for each of the curves from (D) with standard error and logarithmic regression best fit (dotted line).

Since the maximum OD is determined by the ratio of phage to bacteria, and the number of bacteria at a given OD is unlikely to change, we hypothesized that the effect we observed could be due to changes in the number of effective phages present. For example, if the phage population were to lose potency over time, that would have a larger effect on the cultures with lower initial population sizes because they require more time to reach their respective maximum OD. In this scenario, when there are few bacteria in a culture, the rate at which phage encounter the bacteria goes down, causing them to reproduce more slowly. At the same time, if phages lose potency with time, the extra time it takes for the cultures with lower starting bacterial populations to reach the same OD levels of those that started with more bacteria would provide time for the phage to decay. This decay would in turn allow for the bacteria to grow more before the population collapses, causing the maximum OD to increase (Figure 2B).

Phages are typically thought to be highly stable and have even been documented to survive long periods of storage in harsh conditions [5,6], such that the hypothesis that phages were decaying during our experiment was counterintuitive. Nevertheless, we directly tested if the lambda(vir) phage would lose potency in the conditions of our experiment. Similar to the previous experiment, we started with a constant number of phages but this time we waited for different lengths of time before exposing bacteria to the lambda(vir) phage. Specifically, we shook phages in LB media at 37°C for 0, 0.75, 1.5, 3, and 16 hours before adding 10^7 *E. coli*, with each well in a row of a 96-plate representing replicates of a different phage growth time delay (Figure 2C). If phage potency were to decay with time, we would expect that there would be a positive correlation between the length of the delay and the maximum OD.

Our time delay experiment revealed that the phage that shook for longer periods of time resulted in cultures with higher maximum ODs than those of phage that were incubated for shorter time periods (Figure 2D). We observed a logarithmic correlation between maximum OD reached and the delay length before bacteria were introduced to the culture (Figure 2E). This change in maximum OD mirrored the change in maximum OD observed in our first experiment (Figure 2A), supporting our hypothesis that phage decay over time during the experiment, and that this decay causes changes in maximum OD. We note that in all cases the population eventually collapsed, such that the finding that phage potency decays during the course of the experiment could only be made by following the population dynamics, and not just the endpoints.

#### The debris of dead bacteria can significantly affect OD measurements

In addition to the surprising change in maximum OD, we noticed a second unexpected trend in our initial experiment: the final OD at which the culture plateaued changed as a function of the initial population size (Figure 3A). Specifically, the lower the initial population size was, the higher the OD of the plateau (Figure 3B). This was unexpected, because after phage take over a system and the bacterial population collapses, the bacteria left alive would be minimal and below the limit of spectrophotometer detection for the OD to plateau.

**Figure 3.**
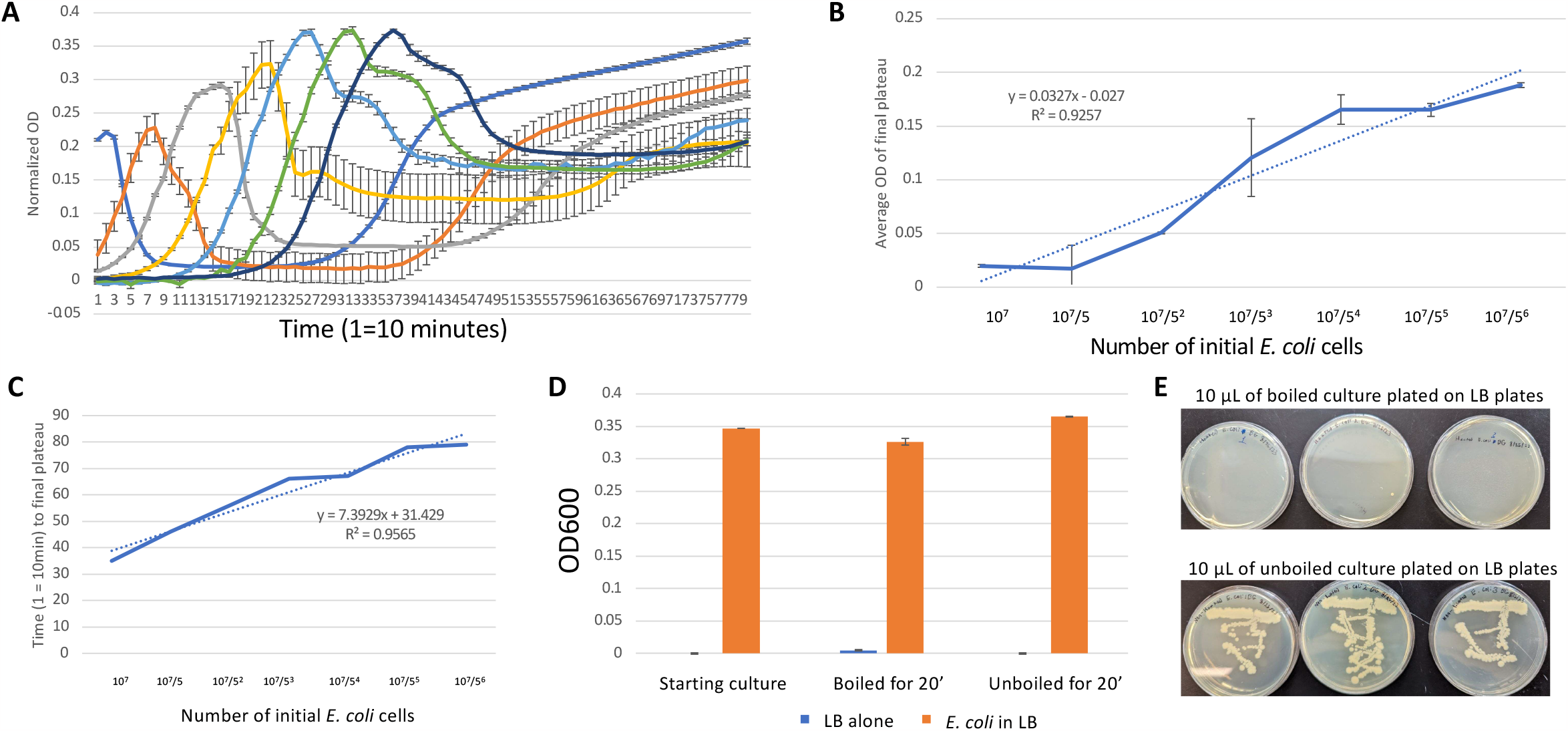
(A) Averages of the replicate wells from each row from the experiment in Figure 1 to focus on the OD of the final plateau. Colors represent the rows illustrated in Figure 1D in descending order from most starting cells to fewest of: dark blue, orange, gray, yellow, light blue, green, navy, brown. (B) Average OD of the lowest value in the final plateau as a function of initial cell number. Error bars reflect standard error of the mean. (C) Time required for each average curve (from A) to reach its final plateau. Dotted line represents a linear regression. (D) Average OD600 values for LB media alone (blue bars, without bacteria) and an LB bacterial culture (orange bars), before treatment (starting culture), after 20’ of boiling (boiled for 20’), and after 20’ at room temperature (unboiled for 20’). Error bars represent standard error of the mean. E. Images of LB plates after 24 hours of incubation that were each plated with 10 μL of boiled culture (top row) or unboiled culture (bottom row).

Examining the time that it took to reach this minimum value of the final OD plateau, we noticed a correlation between the time that it took for a culture to plateau and the final OD value at which the plateau occurred (Figure 3C). This correlation implied that the longer it took to reach the minimum value, the larger that value would be. One thing that similarly changes with respect to time is the accumulated number of bacteria that die as a result of phage infection. Thus, while we had initially assumed that OD measurements reflect the number of living bacteria present, we hypothesized that the buildup of the remains of dead bacteria could potentially explain the change in OD plateau.

To test whether dead *E. coli* would be detected in an optical density measurement, we sought to directly kill bacteria by a method other than phage lysis and measure the resulting impact on OD. To kill the bacteria, we boiled the *E. coli* in LB media for 20 minutes. As positive and negative controls, we also boiled bacteria-free LB media, and compared our results to unboiled samples. We observed only a negligible change in OD of the unboiled LB vs boiled LB (Figure 3D), showing that boiling the LB media itself had a minimal effect on the OD. In contrast, comparing boiled and unboiled *E. coli* cultures revealed that boiling only partially reduced the culture OD, indicating that the debris of the dead bacteria left a significant and measurable effect on OD (Figure 3E). To confirm that the boiling actually killed the *E. coli*, we plated samples from both the boiled and non-boiled groups to determine if viable bacteria were still present. We saw substantial growth from the *E. coli* that were unboiled, and detected no growth whatsoever from the *E. coli* that were boiled (Figure 3F). These results indicate that the remains of dead bacteria can have significant effects on OD measurements.

## Discussion

Here we used high-throughput OD measurements to follow the dynamics of phage-bacterial interactions, using *E. coli* MG1655 and the lytic lambda(vir) phage as a model system. Performing a bacterial cell number dilution series experiment in multiple replicates revealed that the variance in the time required for phage-resistant mutants to emerge increased as initial population size decreased. This conclusion mirrors that of the original experiments performed by Luria and Delbruck [2], but required significantly less time and effort to get the same result. Importantly, however, in addition to replicating expected results, we found that monitoring the dynamics of the system revealed unexpected patterns that inspired additional experiments. Specifically, without following the culture OD throughout the experiment, we would not have noticed other changes that correlate with the starting cell number, such as the number of bacteria that accumulate before the culture crashes before the phages take over or the final OD at which the culture stabilizes after the culture crashes.

We discovered that the changes in bacterial accumulation before culture crash (peak size) could be explained by the decay in phage potency during the experiment. We directly demonstrated that lambda(vir) decrease in potency as a function of time spent shaking in LB media at 37°C. This effect explains the change in peak size because cultures with fewer starting cells take longer to reach the same final cell number, and therefore have fewer active phages present once they reach those levels. This observation was surprising because it goes against the common assumption in the phage field that phage are stable when they are not growing [5,6]. Our results that phage significantly decay under standard bacterial growth conditions have broad implications, as it means that studies that make such an assumption need to account for this decay. This is especially crucial in scenarios where the quantity of phages is significant. In particular, our findings may be significant for phage therapy [5,6], as our results suggest that phages might rapidly decay while in the body before they reach their targets. From a pharmacokinetic perspective, our results suggest that phage therapy dosages should be changed to account for how long the phage needs to be in the body before reaching its target.

Our second observation, that the final culture OD plateau changes as a function of starting cell number, could be explained by the accumulation of the debris of dead bacteria. Traditionally, optical density is thought to reflect the number of living bacteria in a culture. In contrast, our finding demonstrates that dead bacteria still have a significant impact on optical density. Thus, any optical density study with an agent that can cause bacterial death (like phages, antibiotics, or environmental toxins) must take this point into account.

We note that had we followed the end-points of our experiments rather than their dynamics, we would not have been able to see that phage decayed and dead bacterial debris accumulated. This highlights the value of dynamical measurements, as they provide additional information about a system. Historically, the disadvantage of dynamical measurements was their throughput, as it was difficult to follow the dynamics of many cultures in parallel. The advent of high throughput instruments like plate-reading spectrophotometers makes such high-throughput measurements of dynamics feasible. In the future it should be interesting to revisit other classical experiments that were originally performed using end-point analyses to determine if dynamical measurements might reveal new insights.

## Materials and Methods

### Bacterial and phage strains

All experiments were performed using *E. coli* MG1655 bacteria and lambda(vir) phage.

### Media and growth conditions

All experiments were performed using Luria Bertani media (10 g/L tryptone, 5 g/L yeast extract, 10 g/L NaCl in distilled water) shaking (250 rpm) at 37°C.

### Dynamical OD measurements

The dynamics of the culture OD was assayed using 96-well plates in a Tecan Infinite M200 Pro microplate reader at 37°C shaking for 16 hr. Optical density (OD_600_) was measured every 10 min for each well in the plate.

### Bacterial killing experiments

To directly kill bacteria, we boiled the cultures for 10 minutes. We also assessed bacterial viability afterwards by plating 10 μL of the sample on an LB plate.

### Statistics and data analysis

All data analysis were performed in Microsoft Excel. Standard Error of the Mean is used for all error bars. Linear or logarithmic regressions were performed as indicated and the resulting equations and R-squared values reported on the appropriate graph.

## Acknowledgements

We wish to thank several members of the Princeton University Department of Molecular Biology for technical assistance, including Jacob Marogi and Tom Silhavy. We also wish to acknowledge Michael Tang and Stephanie Shoop for assistance in initiating this project. Finally, we thank Ryland Young for helpful discussions about bacteria-phage interactions.

